# Microbial community response and recovery through an aeration-cessation time series in a eutrophic estuary

**DOI:** 10.64898/2026.09.24.754275

**Authors:** William F. Schroer, Yue Zhang, Keith Arora-Williams, Alice Turnham, Lora Harris, Andrew Heyes, Jeremy M. Testa, Laura L. Lapham, Sarah P. Preheim

## Abstract

Eutrophication-driven hypoxia and harmful algal blooms are an expanding threat to water quality in estuaries. Engineered aeration is a technology used in freshwater and, to a lesser extent, estuarine systems to mitigate both hypoxia and algal blooms. However, the efficacy of engineered aeration has not been well studied within estuarine ecosystems and very little is known about the impact of aeration on microbial communities in either estuarine or freshwater environments. Here we present a time series investigation of microbial community response to aeration and cessation of aeration in a eutrophic estuary. Samples for 16S rRNA gene analysis were collected over about a month-long period during which time the aerators transitioned from being “on” to “off” to “on” again. Cessation of aeration selected for the dominance of a eukaryotic algae similar to *Heterosigma akashiwo*. After the resumption of aeration, *H. akashiwo* dominance decreased rapidly. Cessation of aeration also shifted the structure of the entire microbial community, where diversity decreased and the structure of microbial communities at the surface diverged from those from the bottom of the water column. These changes did not occur at non-aerated control sites. Distinct groups of taxa responded to the disruption in a successional pattern through time, with most taxa never returning to their pre-aeration-cessation abundance during the observational period. These findings demonstrate the complex community response to anthropogenic interventions and provide support for bloom suppression through aeration.

**Importance:** Hypoxia and algal blooms represent a severe and growing threat to the economies of coastal communities. Engineered aeration has been well studied as a tool to mitigate hypoxia and algal blooms in freshwater systems, but little is known about its efficacy in estuarine systems or its impact on the whole microbial community. Our results demonstrate that, within a eutrophic estuary, aeration mitigates hypoxia, suppresses algal blooms, and significantly alters the broader microbial community structure. Stopping aeration dramatically restructures microbial community composition, and these microorganisms don’t immediately recover after two weeks of altered oxygen conditions. This work shows that engineered aeration is an effective tool maintain water local quality in small estuaries, but also offers the warning that any cessation, such as from failures or planned outages, can cause large disruptions to microbial communities and may lead to rapid algal bloom formation.

## 1. Introduction

Hypoxia (dissolved oxygen < 2 mg/L [1]) is a rapidly growing threat to estuarine and marine water quality, largely driven by eutrophication associated with excess nutrient loading [2–4]. Hypoxia negatively impacts both biological and chemical processes in estuarine systems by reducing habitat availability for macro-fauna dependent on oxygen for respiration, triggering the release of sediment bound phosphorus [5] that can exacerbate eutrophication, and shifting microbial community function towards the production of noxious (e.g., hydrogen sulfide [6]) and greenhouse gasses (e.g. methane [7]). Addressing excess nutrient runoff represents the root cause of eutrophication-driven hypoxia but is a complex issue that often requires government, community, and corporate cooperation at the watershed scale. This can leave local municipalities and communities searching for other options to manage hypoxia in their local estuaries. One approach to mitigate hypoxia is the use of engineered aeration (hereafter “aeration”). This can take the form of many design solutions. Here, we specifically explore fine bubble diffusion of compressed air through a bubble emitter placed on or above the sediment surface. Fine bubbles reoxygenate the water through a direct gas exchange [8]. Aeration causes abrupt changes to the chemical and physical environments, but it is unclear how these changes will impact the structure of aquatic microbial communities that mediate critical biogeochemical cycles.

Aeration has been found to suppress harmful algal and cyanobacterial blooms (HABs) in freshwater systems. However, the impact of aeration on algae has not been studied in estuarine environments. The rich body of literature covering studies of aeration in freshwater systems (e.g. [9–12]) frequently propose two mechanisms by which aeration may mitigate HABs. These are (1) water column mixing from water-lifting bubbles transports algal cells out of the sunlit photic zone and (2) reduced P availability driven by reduced internal P loading associated with increased oxygen concentrations [9]. Aeration is generally thought to be most effective at reducing cyanobacterial blooms, as these cells are more negatively impacted by mixing than Eukaryotic algae which have a wider range of optimal light intensity [9]. The extensive body of literature on this topic, however, paints a more complex picture. Reviews of aeration studies in freshwater lakes by Visser et al [9] and Pastorok et al [10] show that artificial mixing of the water column, via aeration and other mechanisms, was only effective at reducing cyanobacterial populations in 57% of reported cases (21 out of 37) [9, 10]. It appears that the efficacy of aeration mixing is highly variable on a case-by-case basis and that, even within the same lake, conflicting outcomes have been observed from repeated investigation [9, 10]. Similarly, mixed efficacy has been shown in more recent publications. Lerminiaux et al (2026) [11] compared algal bloom dynamics between five aerated and five unaerated farm ponds and found that aeration had no significant effect on total chlorophyll-a or cyano-toxin concentration. However, Naderi, et al. 2022 [12] found that, in a reservoir, over 6 days of aeration, cyanobacterial populations decreased by ∼50% and green algal populations decreased by ∼20%. This high variability of algal and cyanobacterial response to aeration suggests that there are factors influencing the outcome that have not yet been identified.

One possible explanatory factor for variable results of aeration on phytoplankton blooms could be the influence of the broader microbial community. Algal cells exist within diverse microbial communities and participate in numerous mutualistic and antagonistic interactions with bacterial co-members [13–16]. Aeration driven disruptions of the broader microbial community may generate feedbacks, positive or negative, for algal taxa. Though microbial communities are key drivers of biogeochemical processes, microbial community response to aeration is rarely measured. One study that tracked microbial community response to water-lifting aeration [17] found differences in community composition between control and aerated sites, mainly focusing on differences associated with physical (temperature) and chemical (oxygen, nitrogen, and carbon) changes caused by aeration. They also found that aeration increased diversity relative to a control site. While phylum-level changes were modest over time and with aeration [17], it is not clear whether more dramatic changes occur at finer taxonomic resolution in response to aeration. Elucidating the microbial community’s response to the cessation and re-establishment of aeration at higher taxonomic resolution can provide insight into which taxa are most sensitive to the impacts of aeration and resulting changes in oxygen availability and redox conditions.

Here we monitor the structure of the microbial community through an aeration – cessation time series in a eutrophic estuary. The goals of this study are to determine the impact of aeration and aeration cessation in an estuarine ecosystem on (1) photosynthetic microorganisms, including potential HABs and (2) bacterial community structure. During July, 2019, we collected a time series of samples for 16S rRNA gene analysis from Rock Creek, a tidal tributary estuary of Chesapeake Bay that has been seasonally aerated since 1988 to mitigate seasonal hypoxia [18–20]. Samples were collected before, during, and after a planned shutdown of the aerators. The rapid and dramatic changes in community structure, including algal bloom development, provides insights into the stabilizing role aeration can play in eutrophic estuaries. Our findings show that even after decades of aeration, a brief cessation in operation can trigger harmful algal blooms. This supports the role of aeration addressing the acute threats of hypoxia and HABs, but not resolving the underlying drivers, such as eutrophication.

## 2. Results and Discussion

### 2.1 Cessation of aeration triggered an algal bloom

The head waters of Rock Creek, Pasadena, MD, USA are typically aerated for 12 hours daily during the summer months to mitigate the threat of hypoxia (standard operation designated as “on”) (Methods, Fig. 1) [5, 7, 19]. This aeration is currently accomplished by pumping compressed air through emitters that produce ∼3 mm bubbles capable of directly oxygenating the water through gas exchange as well as overturning the water column [5, 7]. During July of 2019 the aerators were turned off for a period of 12 days (“off”), after which point they were run on an altered schedule for four days (“altered”), before returning to standard operation (“on”) (Table S1, Fig. 1b). To elucidate the impacts of aeration and aeration cessation on the structure of the microbial community, 115 samples were collected for 16S rRNA gene sequencing prior to, during, and after these manipulations. Samples were collected from two stations immediately adjacent to the aerators (RC1, RC2) and two stations (RC7, RC9) 1-2 km distant from the aerators (Fig. 1). The samples from the non-aerated stations RC7 and RC9 represent valuable reference points for comparison with aerated sites, but they are not true control samples. Hydrologic connectivity and tidal conditions within the Rock Creek estuary prevent RC7 and RC9 from being completely independent of RC1 and RC2. Additionally, RC7 and RC9 are down bay of RC1 and RC2, where the estuary is wider and there is greater fetch and opportunity for mixing with the main basin of the Patapsco River estuary near its boundary with Chesapeake Bay.

**Figure 1.**
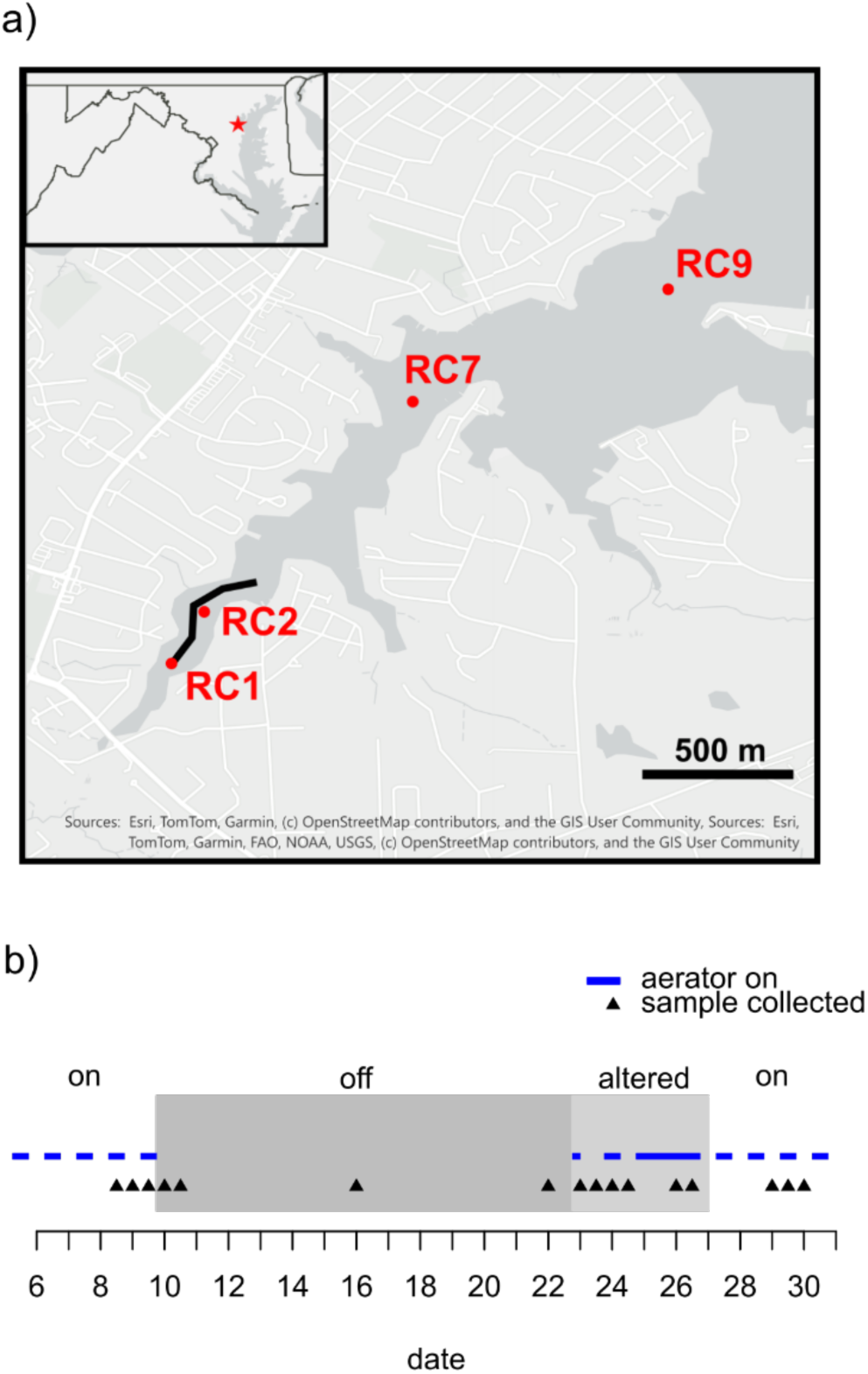
(a) Map of the Rock Creek tributary in Pasadena, Maryland, USA. Water samples were taken at two sites (RC1 and RC2) adjacent to the aerators (marked by the black line) and two sites downstream from the aerators (RC7 and RC9). (b) Timeline of aeration and sampling in July 2019. Blue dashed bars indicate times aerators were running. Gray and light gray boxes indicated the time periods considered “off” and “altered”, respectively. Black triangles mark the dates at which water samples were collected.

The most dramatic response to the cessation of aeration was the increase of chloroplast 16S rRNA gene sequences in the dataset. Traditionally, the 18S rRNA gene is used for eukaryotic algal analysis, however chloroplast 16S rRNA gene abundance is an effective alternative [21] and has been shown to be more sensitive than 18S rRNA genes for identifying co-occurrence patterns between phytoplankton and heterotrophic bacteria in a marine time series [22]. Phytoplankton amplicon sequence variants (ASVs), including both cyanobacteria and chloroplast genes, were generally prevalent in surface samples throughout the experiment (Fig. 2). Prior to cessation of aeration, *Synechococcus* was the dominant phytoplankton taxa representing a read fraction of about 0.2 (20%). Within 6 days of cessation, at RC2, there is a spike in the abundance of a single ASV with 97% identity to the chloroplast of *Heterosigma akashiwo*. *H. akashiwo* is a mixotrophic Raphidophyte alga that has been shown to feed on heterotrophic bacteria and *Synechococcus* [23], and causes harmful algal blooms in coastal water bodies around the world [24, 25]. Within 12 days, *H. akashiwo* was the dominant taxa in surface water samples at both RC1 and RC2, at its peak representing a read fraction of 0.8 (80%) of all reads in those samples (Fig. 2). The bloom reached its peak and began to terminate during the period of “altered” aeration, with *H. akashiwo* rapidly falling to ∼10% abundance during the return to typical aeration conditions. Quantitative PCR data (Fig. S1) show that the peak of the bloom corresponded with a peak in 16S rRNA gene copies per ml. This indicates that the increase in *H. akashiwo* abundance is indicative of a true increase in biomass, as opposed to being an artifact of relative abundance. This is also supported by total chlorophyl concentrations (Fig. 2, [5]).

**Figure 2.**
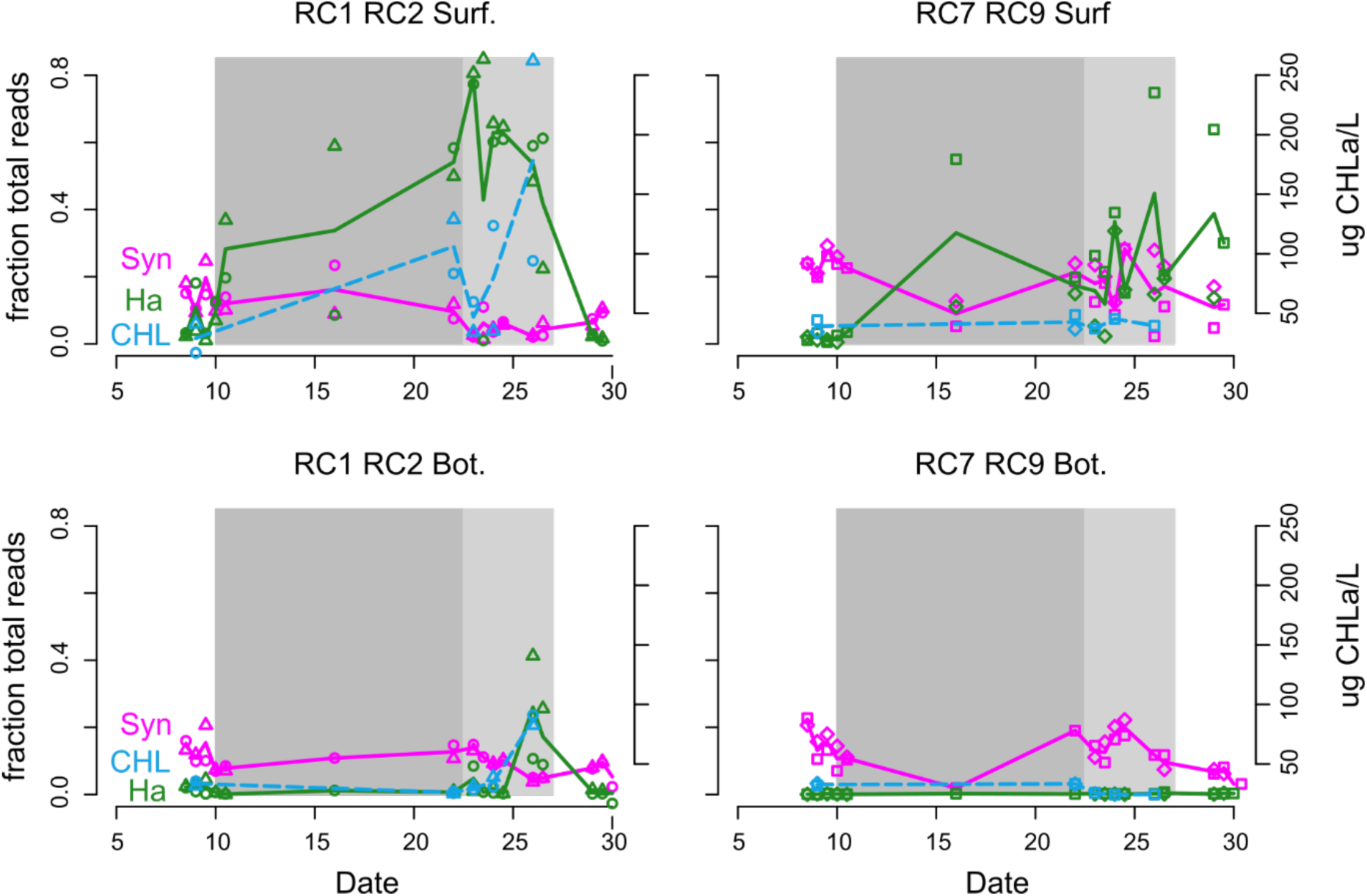
Phytoplankton relative abundance though time. Data are plotted by station group: aerated (RC1 RC2) vs non-aerated (RC7 RC9), and depth in water column: surface (Surf.) vs bottom (Bot.). Magenta indicates 16S rRNA gene read abundance of *Synechococcus*, green indicates 16S rRNA gene read abundance of *H. akashiwo* chloroplasts, blue (and dashed line) indicates total Chlorophyl concentration μg/L. Point shape indicates sample station: RC1 (circle), RC2 (triangle), RC7 (square), RC 9 (diamond). Lines show mean value at each time point. Gray and light gray boxes indicated the time periods considered “off” and “altered”, respectively. Chlorophyl data originally published in Testa et al 2025 [5].

To confirm the identity of the *H. akashiwo* relative in our bloom, we analyzed two shotgun metagenomic samples that were collected during the peak of the bloom. We were able to identify a contig that contained a gene with 97.3% identity to the *H. akashiwo* 18S rRNA gene, further confirming the presence of an alga closely related to *H. akashiwo*. In these samples, the putative *H. akashiwo* contigs had per base coverage of 500-800 times greater than median per base coverage. Additionally, we directly mapped metagenome reads from the two surface samples to the type strain CCMP452 genome (Bowtie2, [26]). However, only 1% to 2% of total reads, respectively, were successfully mapped. The reads mapped unevenly to the CCMP452 genome with contig coverage ranging from 27% to 0% (median 2%). Bowtie2, with default settings, is a generally conservative read mapper [26], the low read map rate likely indicates genetic divergence between the alga in Rock Creek and *H. akashiwo* type strain.

A follow-up field campaign was performed in July of 2022 with a more limited sampling regime. The aerators were turn off on 7/25/22 and a HAB rapidly formed, this time causing a fish kill. On 7/27/22, during the peak of the bloom, staff from the Maryland Department of Environment investigated the reported fish kill. They observed 800 dead fish and attributed the event to a bloom of the dinoflagellate *Levanderina fissa* (aka *Gymnodinium*), which was present at 2.2e4 cells ml^-1^ (Table S2). This cell density represents a severe bloom, previously documented *Gymnodinium* blooms reach cell densities of ∼1e2 to ∼1e4 cells ml^-1^ [27–29]. In response to the speed and severity of negative impacts on the local community, the aerators were returned to operation on 7/28/22, prior to the scheduled date of 8/1/22. Due to the altered aeration schedule, samples for DNA analysis could not be collected during the peak of the bloom, but shortly after on 7/30/22. Analysis of the 16S rRNA gene sequences from samples collected before and after the fish kill event were consistent with an increase in chloroplast abundance (Fig. S2), though there are no published sequences of the *L. fissa* chloroplast 16S rRNA gene against which we can compare. As occurred in the 2019 experiment, *Synechococcus* was the dominant phytoplankton prior to the cessation of aeration and cessation prompted a regime change in favor of mixotrophic eukaryotic phytoplankton. This replication pattern suggests the cessation of aeration generates conditions that promote the growth of mixotrophic Eukaryotic algae more generally, not just *H. akashiwo*. Also, it appears that *Synechococcus* is well adapted to the conditions that exist under the standard aeration regime at Rock Creek, though populations are maintained at low (i.e. non-bloom) levels.

Our data demonstrate that, within this system, aeration inhibits bloom formation and cessation of aeration triggers a bloom. The mechanism of this is not known but we propose three hypotheses. First, the physical action of the aerators themselves could create selective pressure against these eukaryotic algae. Density profiles suggest that the fine bubble aeration system in Rock Creek can mix the water column (Fig. 3b,c). In freshwater systems, water column mixing is thought to preferentially benefit negatively buoyant algae while inhibiting positively buoyant *Microcystis* [9]. This would be mechanistically consistent with our observations as the negatively buoyant *Synechococcus* [30] is more abundant prior to cessation of aeration. *H. akashiwo* and *L. fissa* are positively buoyant [31, 32] and are inhibited by aeration. Additionally, dinoflagellates can be sensitive to turbulence and shear stress [33]. The turbulence produced by fine bubble plumes could be sufficient to inhibit growth through that mechanism [34].

**Figure 3.**
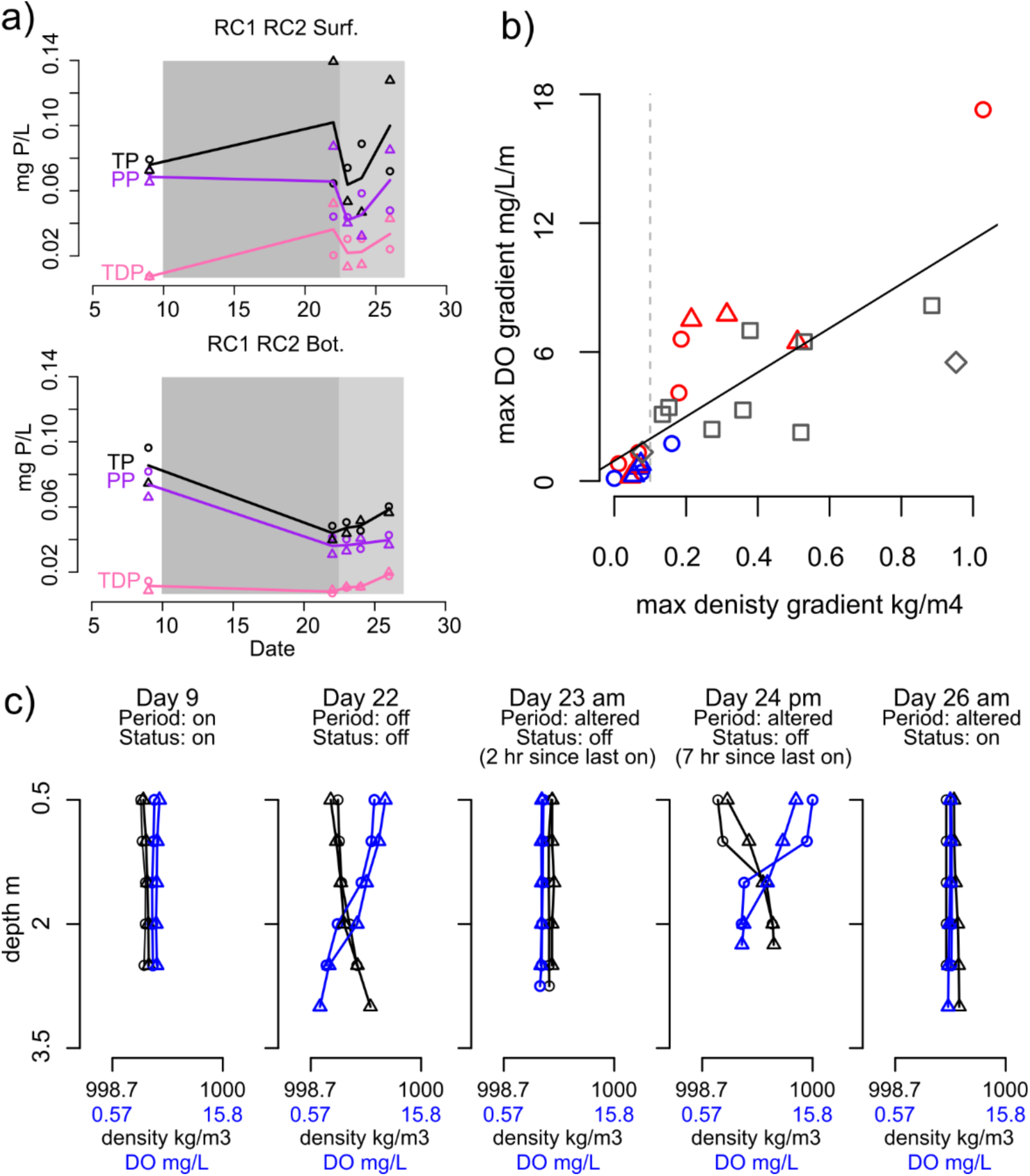
(a) Water column concentrations of total P (TP, black), particulate P (PP, purple), and total dissolved P (TDP, pink) in surface and bottom waters at aerated stations RC1 and RC2. Gray and light gray boxes indicated the time periods considered “off” and “altered”, respectively. (b) Scatter plot of maximum density gradient and maximum dissolved oxygen gradient from all depth profiles. Red points indicate aerated sites with aerators off, blue points indicate aerated sites with aerators on, gray points indicate non-aerated sites. Gray dashed line indicates the 0.1 kg/m^4^ threshold that is used to define a stratified water column in Chesapeake Bay. Solid black line gives the linear regression trendline (R^2^ = 0.60, p=2×10^-6^). (c) Depth profiles of water density (black) and dissolved oxygen (blue) at aerated sites RC1 and RC2 on selected dates. For all plots point shape indicates sample station: RC1 (circle), RC2 (triangle), RC7 (square), RC 9 (diamond). Phosphorus data originally published in Testa et al 2025 [5], profile data originally published in Lapham et al 2022 [7].

The second hypothesized mechanism by which aeration may suppresses algal blooms is through phosphorous release from the sediment, which limits growth of eukaryotic phytoplankton (e.g. [9]). While P release from sediments during anoxia was observed during previous experiments in 2012, 2016 and 2018 at this site when aeration was manipulated, the P release was substantially diminished in 2019 [5]. Additionally, there was no observed accumulation of total dissolved P (TDP) in bottom waters at aerated sites during the off phase (Fig. 3a).

Finally, we hypothesize the eukaryotic algae could benefit from P release from dead or dying bacteria not adapted to the new conditions or consume them directly [35]. Based on qPCR, the total number of 16S rRNA gene copies drops after the cessation of aeration, consistent with microbial community die-off (Fig. S1). Previously published nutrient data [5], show that prior to cessation of aeration almost all P in surface waters was in the form of particulate phosphorus (PP), which includes cellular biomass (Fig. 3a). At RC1, during cessation there was an approximately proportionate decrease in PP and increase in total dissolved P (TDP), consistent with a liberation of P from biomass (Fig. 3a, hollow circles). At RC2 both PP and TDP increased, which would be consistent with an input of P from an external source (Fig. 3a, hollow triangles), though it is unclear what this source may be. These hypotheses may also not be mutually exclusive, multiple mechanisms may have contributed to bloom formation and termination.

### 2.2 Dynamic community response to cessation of aeration

Beyond the algal bloom, cessation of aeration represented a perturbation to the whole microbial community. Continuous oxygen monitoring data collected during the duration of this experiment from bottom waters near station RC2 have been previously published [5], as well as oxygen and density profiles (Fig. 3c, [7]). Continuous monitoring data show that less than two days after cessation of aeration, bottom waters fell below the hypoxic threshold of 2 mg/L, dissolved oxygen remained below or near this threshold for the duration of the “off” period [5]. Density profiles, calculated based on temperature and salinity [36], identify periods of water column stratification (Fig. 3c). Peak density gradients in this study ranged from 0 to 1.03 kg/m^4^ (Fig. 3b). In Chesapeake Bay a peak density gradient of 0.1 kg/m^4^ is considered the threshold used to define a stratified water column [36]. The water column was less likely to become density stratified during periods when aerators were actively running than when they were off (Fig. 3b). The magnitude of the density gradient was positively correlated with the oxygen gradient (Pearson’s r = 0.78, p = 2E-6) (Fig. 3b).

Consistent with the observed chemical and physical stratification of the water column, cessation of aeration led to divergence in the structure of surface and bottom water microbial communities. Immediately following the cessation of aeration, the Weighted UniFrac distance, a measure of community dissimilarity, between paired surface and bottom samples began to increase at the aerated sites (Fig. 4a). The Weighted UniFrac distance remained high throughout the “off” period and began a decreasing trend during the “altered” period, finally returning to near their initial low values during the return to normal aeration conditions. At these sites, the dissimilarity between surface and bottom communities was significantly higher during the “off” and altered period than the “on” periods (Wilcox p=7e-4). These results are consistent with the development of a bloom at the surface that did not extend to the bottom. This may also reflect new, distinct niches in each layer of the water column. At the non-aerated sites, Weighted UniFrac dissimilarities between surface and bottom samples showed a general increasing trend through time, although they were not significantly different during the “on” versus “off/altered” periods (Fig 4a, Wilcox p>0.05).

**Figure 4.**
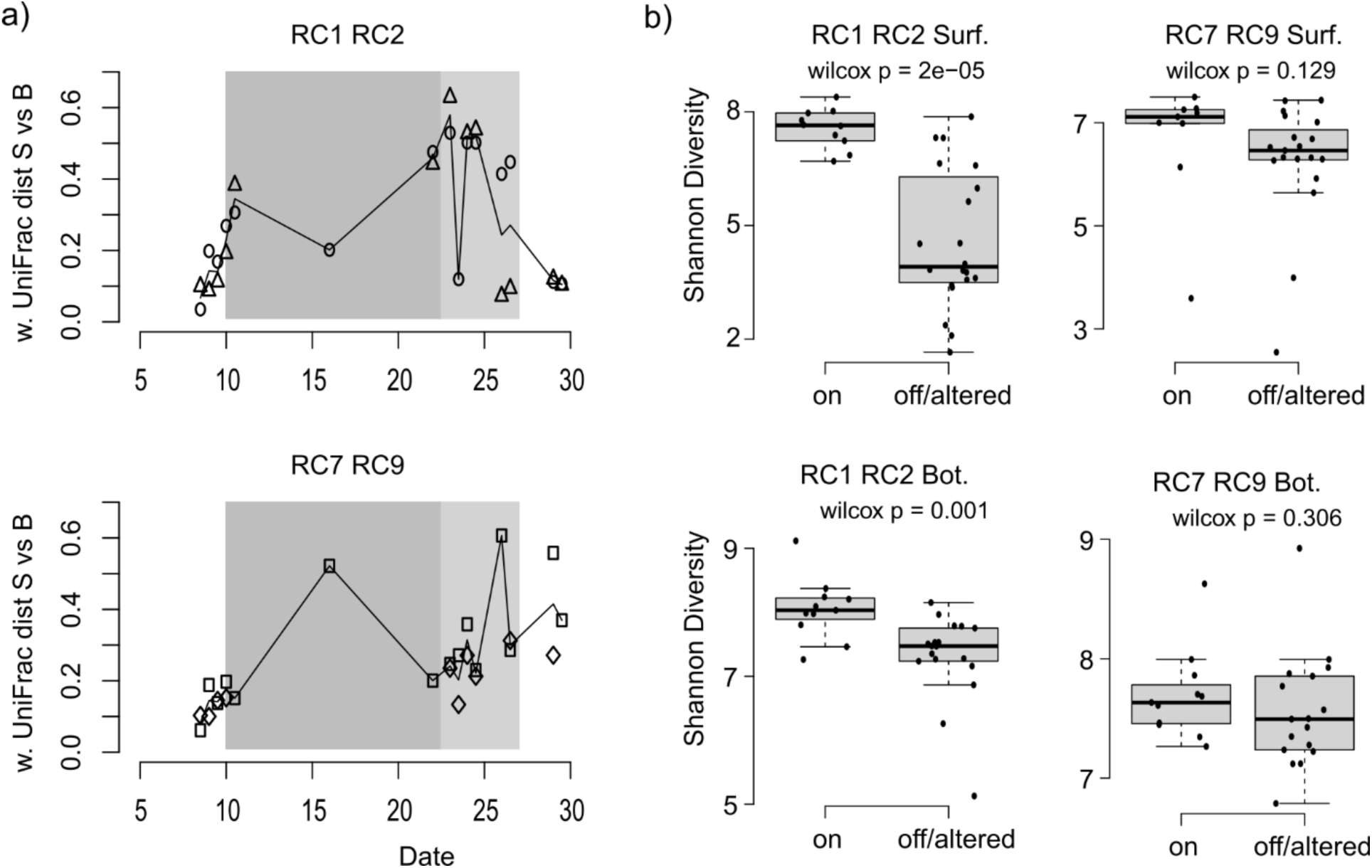
(a) Weighted UniFrac distance between paired surface and bottom samples at all stations. Point shape indicates sample station: RC1 (circle), RC2 (triangle), RC7 (square), RC 9 (diamond). Lines show mean value at each time point. Gray and light gray boxes indicated the time periods considered “off” and “altered”, respectively. (b) Box and whisker plots showing Shannons diversity index at each station group and depth during the “on” periods and the “off”/”altered” periods. Solid black circles show all data points from which the box and whiskers were determined. Also displayed on each plot is the p value from a non-parametric Wilcox test comparing Shannon index between the two time groups.

Beyond the indication of community stratification, alpha diversity metrics show that cessation of aeration reduced community diversity, which recovered upon resumption of aeration. In surface water samples, Shannon’s Diversity Index was significantly lower during the “off”/“altered” periods than during the “on” periods (Fig. 4b). This diversity loss in surface waters is driven by the *H. akashiwo* bloom, in which a single ASV represented more than 70% of all reads in several samples. However, bottom waters, which were largely isolated from the *H. akashiwo* bloom, also experienced significant, though less dramatic, reductions in diversity during “off” and “altered” periods (Fig. 4b). These reductions in diversity are likely a response to the disturbance caused by the rapid change in the chemical environment. These results are consistent with the canonical pulse disturbance-recovery patterns that have been demonstrated across various ecosystems [37, 38], with diversity decreasing in response to the disturbance and then recovering to stable state or alternative stable state. Diversity metrics at the unaerated stations did not change significantly during the period of aeration (Fig. 4b).

Though alpha diversity metrics indicate a recovery of diversity with the return to standard aeration conditions, the composition of the community that emerged post disturbance was distinct from that of the pre-disturbance community. NMDS and Permanova based on Weighted UniFrac distance shows that at all stations the community composition post bloom significantly differs from pre-bloom (Fig. 5, Fig S2). These NMDS plots (Fig. 5, stress: 0.089) show a clear progression of community composition through time at all sites, that can be visualized as the distance between means of samples collected within distinct time periods: “pre-cessation”, “off”, “altered”, and “post-cessation”. At the aerated sites the total distance between time period means was 0.72 and 0.24, for surface and bottom samples, respectively (sum of length of arrows in Fig. 5). These changes are likely driven by the bloom in surface samples and the shift to hypoxia in bottom samples. The surface samples at the aerated sites show a progression of disturbance and recovery with respect to NDMS axis 1, and continuous shift in community towards lower values on NMDS axis 2. The bottom water samples predominantly show the shift to lower values along NMDS axis 2 (Fig. 5). As a comparison, at the non-aerated sites, though there is significant change in community composition through time (Fig. S3), the magnitude of these changes is modest, with total distance between time period means of 0.18 and 0.12, for surface and bottom samples, respectively. An investigation of taxa loadings that influence the NDMS ordination show that axis one is dominated by a gradient of actinobacteria to chloroplasts (Fig. S4). Axis two is characterized by a diverse group of classes driving positive values, while negative values are influenced by Acidimicrobiia and Thermoleophilia (Fig. S4). These shifts in community structure are likely driven by some combination of natural community drift and hydrologic connectivity with the aerated sites. Community diversity metrices indicate meaningful changes in community structure through time but they do not provide information about the taxa that are involved in these shifts.

**Figure 5.**
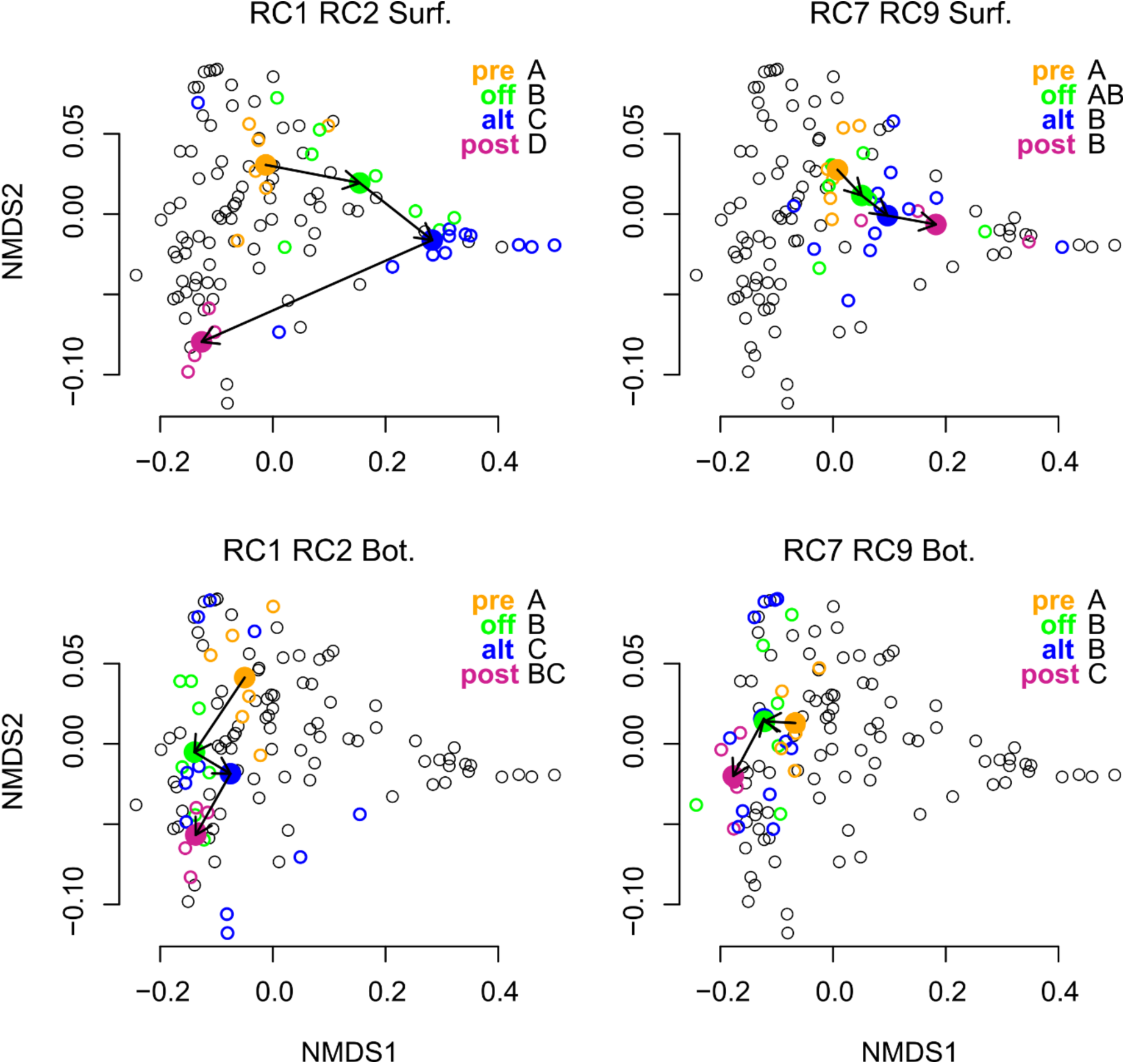
NMDS plots based on Weighted UniFrac distance samples (stress = 0.089). Values for all samples are displayed across all panels (black open circles). Each panel highlights the samples for a given station group and depth, listed in the plot title. Within each panel, samples are colored by the time period in which they were collected. The mean of each time period displayed as a solid circle. Arrows indicate the “movement” of mean through time. Significance codes (A, B, C, D) indicate significant differences in community structure between groups (Permanova, adjusted p<0.05).

In bottom waters, shifts in community structure are not directly explained by changing community metabolic function in response to redox conditions, but some metabolic processes show trends that correspond to measured chemical conditions. Metabolic function was predicted based on 16S rRNA gene taxonomy. Surprisingly, these data do not show a shift away from aerobic metabolism and towards anaerobic metabolism during the cessation of aeration (Fig 6a, b). However, there are examples of specific predicted metabolic functions whose abundance patterns correspond to changes in the chemical environment. At the aerated stations, taxa predicted to perform methanotrophy peak in abundance on the 22^nd^, during the off period (Fig. 6c). Dissolved methane concentrations measured on the same date were approximately three times greater than concentration measured prior to cessation of aeration (Fig 6e, [7]). The high methane concentration would provide ample substrate to stimulate the growth of methanotrophic populations. Further, taxa predicted to perform nitrate denitrification increased in abundance immediately after the cessation of aeration and remained elevated during the off period (Fig 6d). This corresponded with a decrease in total nitrogen that was driven by the loss of nitrate (Fig 6f, [5]), consistent with N removal from the system through denitrification. Together these selected examples support a shift in microbial community structure and function towards denitrification and methanotrophy. This combination of metabolisms is somewhat surprising as denitrification is an anaerobic process [39] while water column methanotrophy is generally an aerobic process [7]. The co-occurrence of these two metabolisms could be explained by small scale spatial and temporal variation in DO, as continuous monitoring data [5] and depth profiles (Fig 3a) show conditions fluctuate between fully anaerobic and hypoxic. In other marine systems co-occurrence of aerobic and anaerobic pathways have been explained by cryptic oxygen fluxes [39], which may also occur in this system.

**Figure 6.**
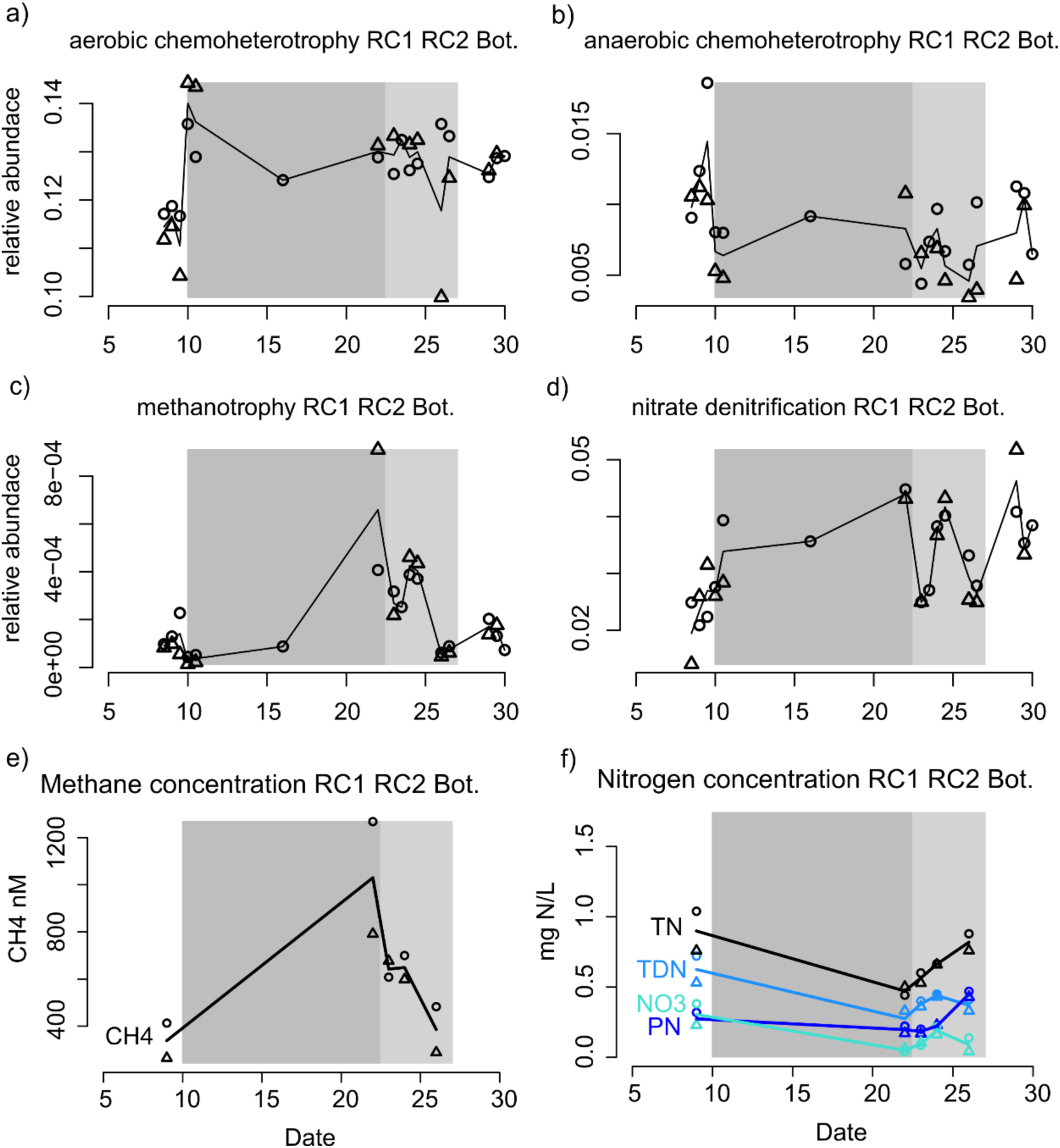
Abundance of predicted of microbial functions from Faprotax2, metabolism listed in figure titles (a-d). Water column methane concentration (e). Water column total N concentration (TN, black), particulate N concentration (PN, blue), and total dissolved N concentration (TDN, light blue) (f). For all plots point shape indicates sample station: RC1 (circle), RC2 (triangle), RC7 (square), RC 9 (diamond). Gray and light gray boxes indicated the time periods considered “off” and “altered”, respectively. Nitrogen data originally published in Testa et al 2025 [5], methane data originally published in Lapham et al 2022 [7].

### 2.3 Patterns of taxa abundance group into six temporal patterns

Tracking abundance at high taxonomic resolution revealed distinct response patterns to the aeration cessation. To assess trends in abundance at the ASV level, ASVs were clustered into modules of correlated abundance [40]. Clustering was performed at each station group and depth independently, for example samples from stations RC1 and RC2 surface were clustered independently of RC1 and RC2 bottom. This analysis produced between 13 and 16 modules at each station-depth group. These modules were then grouped into supermodules based on the time period in which they peaked: “pre-disturbance”, “off”, “altered”, “post-disturbance”, “recovery”, and “no-trend”. Each super module had a distinct taxonomic composition. The use of super modules facilitates identification of specific taxa that follow various patterns of disturbance response.

At aerated sites, 18% - 23% of reads belong to taxa that are present at high abundance prior to the cessation of aeration, rapidly decline in abundance at the onset of aeration and never return to their starting abundance. These are the ‘losers’ of the disturbance and regime shift and include members of betaproteobacteria, *Synechococcus*, betaproteobacteria, actinobacteria, and some Eukaryotic phytoplankton (Fig. 7, S5).

**Figure 7.**
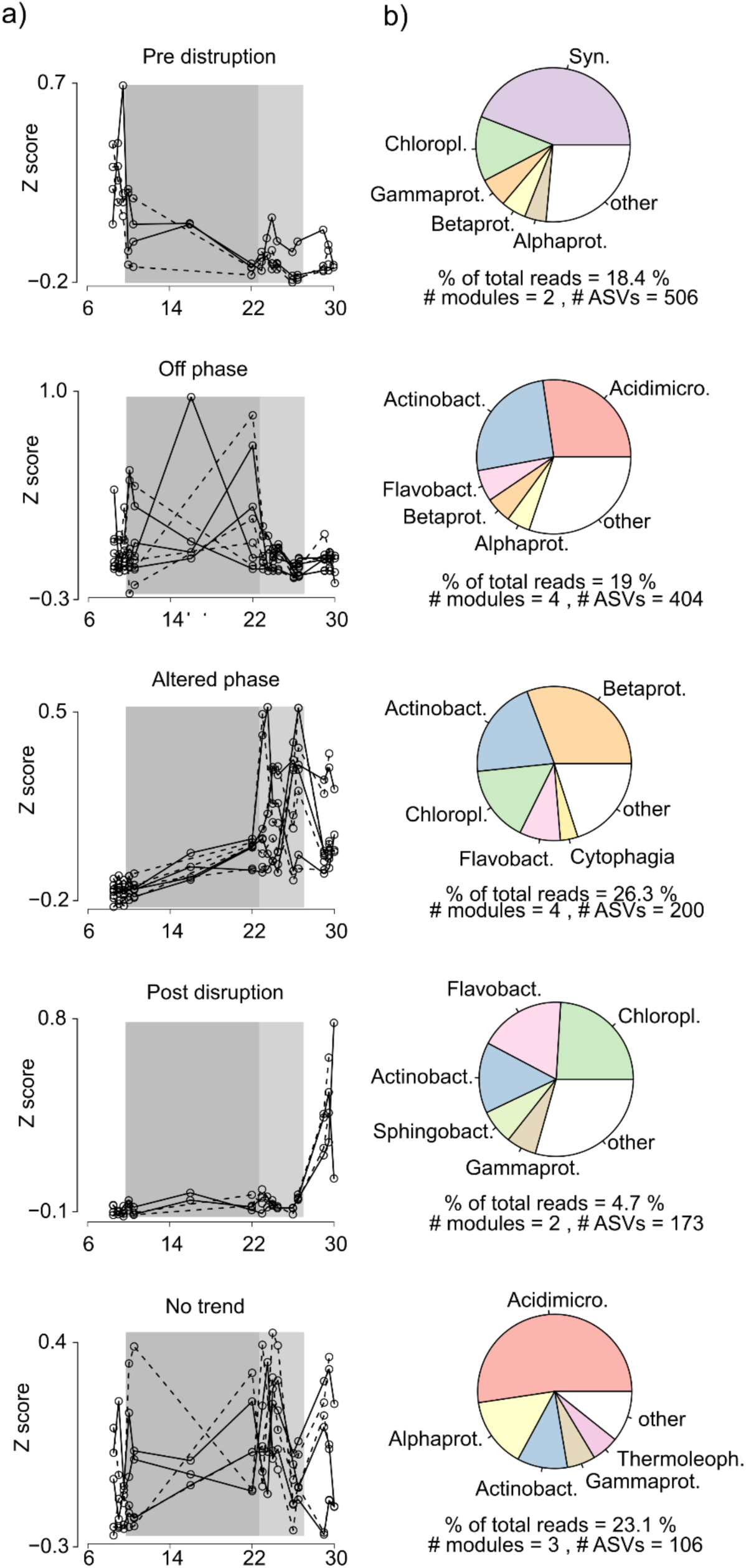
Module data from ASV level WCGNA cluster analysis in bottom samples at aerated stations RC1 and RC2. Plots in panel (a) show Eigengene value through time, a dimensionless value that can be considered a weighted average of ASV Z-score abundance. Each plot depicts the modules that have been assigned to a given super module, listed in the plot title. Solid lines are RC1, Dashed lines are RC2. Gray and light gray boxes indicated the time periods considered “off” and “altered”, respectively. Panel (b) shows pie plots that provide the weighted abundance of all ASVs represented by each given super module. Also in panel (b), is information about the total number of modules in each super module, the number of ASVs represented by those modules, and the total fraction of total reads that map those ASVs.

Two super modules include members that take advantage of the cessation of aeration, those that peak during the “off” and “altered” periods. In surface samples, taxa in these super modules represent ∼54% of total reads and are dominated by phytoplankton. The dominant class of the “off” group is *Synechococcus* (Fig. 7, S5). These *Synechococcus* ASVs, however, are distinct from those in the “pre” super module suggesting niche separation between related strains, this pattern of related ASV’s demonstrating differing abundance patterns is common within this dataset. Proceeding the relatively modest peak of *Synechococcus* is a much larger bloom of *H. akashiwo* related chloroplasts that peaks in the “altered” super module, as discussed in depth above.

As previously discussed, during the “off” and “altered” periods the composition of surface and bottom communities at the aerated sites diverged (Fig. 7, S5). The super modules that peaked during this period in the bottom samples include common anaerobes Actinobacteria and Acidimicrobiia. Though members of both classes are frequently observed in Chesapeake Bay and other aquatic environments [41–43], most research on these taxa comes from soil and sediment systems [44, 45]. Acidimicrobiia are frequently anaerobes and may be capable of complex sulfur and carbon cycling, including carbon fixation [45, 46]. Actinobacteria are known for their production of bioactive secondary metabolites and capacity to degrade complex organic compounds [44]. These taxa are likely important to the function of the Rock Creek and other estuarine microbiome and should be a target of future investigation.

Two super modules increase with the resumption of the standard aeration regime, these are “recovery” and “post-disturbance” (Fig. 7, S5). The recovery super module was only found in the surface water samples at Stations RC1 and RC2. Taxa in this super module represent only 5.7% of total reads, the dominant were members of Actinobacteria and Acidimicrobiia. The “post-disturbance” super module represents taxa that seem the be the ‘winners’, as they only gain abundance during the resumption of normal aeration. In surface samples this module represented 13.4% of total reads and was also largely comprised of Actinobacteria and Acidimicrobiia. As was previously noted for *Synechococcus* ASVs, members of class Acidimicrobiia “off” and “altered” super modules in bottom waters were different from those in the “post-disturbance” super module, indicating successional dynamics between Acidimicrobiia strains.

In bottom samples, the “post-disturbance” super module represented just 4.7% of total reads and the two most abundant taxa groups were Eukaryotic phytoplankton and Flavobacteria. The phytoplankton signal could indicate sinking or mixing biomass coinciding with bloom termination. Flavobacteria have been shown to be involved in the breakdown of algal produced biopolymers, which may explain their increase with the bloom termination signal [47]. The strength of the ASV correlation module approach is the ability to identify divergent patterns between related strains that would be masked by grouping at higher taxonomic resolution (Fig. S6).

ASV modules from the unaerated sites did fall into these same super group patterns, though in both surface and bottom samples a greater portion of total reads (31% and 35%) were represented by the “no-trend” super module than in samples from aerated sites (0% and 23%, Fig S3).

## 3. Conclusions

Engineered aeration is a tool that is widely used to mitigate water quality issues associated with harmful algal blooms and the risks of hypoxia in eutrophic freshwater systems. Here we expand on the existing body of literature by demonstrating the efficacy of engineered aeration in an estuarine system and by monitoring the impact of aeration on the complete microbial community. In Rock Creek where there has been a long-standing aeration regime, the cessation of aeration during the summer represents a substantial disruption to the aquatic microbial community. The cessation of aeration created both physical and chemical changes, which led to a divergence between the surface and bottom water microbial communities. It also led to the formation of an algal bloom in surface waters, as well as a loss of community diversity and shift in community structure throughout the water column. With the resumption of aeration, the algal bloom terminated, for which we have proposed three possible mechanisms. Cessation of aeration three years later triggered an algal bloom with similar dynamics in which *Synechococcus* was the dominant phytoplankton pre-cessation. Rapidly after cessation of aeration, a bloom of mixotrophic dinoflagellates occurred and the resumption of aeration terminated the bloom. Notably, though our results demonstrate that aeration is a valuable tool mitigate the acute threats of eutrophication and hypoxia, they also show that multi-decadal aeration use does not address the underlying drivers of eutrophication and harmful algal blooms.

## 4. Methods

### 4.1 Study site and sampling schedule

Our field site, Rock Creek, Pasadena, MD, USA (Fig. 1), is a tributary estuary of Chesapeake Bay. Its watershed covers 2523-acres of largely residential land. Rock Creek has a long history of eutrophication, harmful algal blooms, hypoxia, and fishkills [18–20]. Aerators were installed in 1988 to mitigate chronic hypoxia and harmful algal bloom formation [18–20]. These initial aerators pumped compressed atmospheric air through a series of coarse diffusers creating large bubbles (∼20 mm) that de-stratify the water column, bringing water to the surface where it can be oxygenated by exchange with the atmosphere. In the spring of 2019, a new aeration system was installed that consist of two 213 m long diffusers that produce smaller (∼3 mm) bubbles these smaller bubbles have greater surface area, allowing for greater direct gas exchange with the water, while still maintaining the capacity to mix the water column [5, 7]. Our main field campaign was conducted in July 2019. During this period, the aerator flow regime was changed from normal operation (“on”) to a period when the aerators were completely off (“off”), then run on an altered aeration schedule (“altered”) before returning to normal aeration (“on”). An overview of this process as well as dates of sample collection are given in Fig. 1 and Table S1.

Samples were collected from 4 sites within Rock Creek (Fig. 1). Two sites, RC1 and RC2, were in the immediate vicinity of the aerators. Two sites, RC7 and RC9, were down bay of the aerators by ∼1 km and 2 km, respectively. Water samples were collected from the surface (∼0.5m below surface) and bottom (∼0.5 m above sediment) of the water column. The water depth was approximately 2.5 meters at RC1 and RC2 and 3.5 meters at RC7 and RC9.

An abbreviated campaign was conducted in July 2022 to determine if the main findings of the 2019 investigation could be replicated. In this campaign Aerators were turned off from 7/25/22 to 7/28/22 and samples were collected on 7/23/22 and 7/30/22.

### 4.2 Water sample collection, storage, and DNA extraction

Water samples were collected in 50 mL sterile centrifuge tubes using a peristaltic pump flushed with 2 volumes of water before sample collection. After collection, water samples were immediately placed onto dry ice and stored at -80 °C for up to eight months. As negative controls, start and end blanks were collected by passing samples of sterile MilliQ water though the peristaltic pump tubing at the beginning and end of each sample collection date. To extract DNA, approximately 50 mL of each water sample was filtered through a 0.22 μm hydrophilic polyethersulfone filter (Millipore Sigma). Each filter membrane was added into a bead beating tube to extract DNA using a DNeasy PowerSoil Pro kit (QIAGEN) following the manufacturer’s protocol. DNA extraction negative controls were run by adding no sample to extraction bead tube.

### 4.3 16S rRNA gene library preparation and sequencing

For the 2019 samples,16S rRNA gene libraries were generated using a previously described two-step amplification method [42, 48]. Briefly, all PCR reactions were performed using Phusion HF polymerase (New England Biolabs, Ipswich, MA). In the first step, the V4 region of the 16S rRNA gene used a pair of universal primers targeting U515F and E786 positions plus adapters for the second step reaction (PE16S-V4-U515-F, 5′-ACACG ACGCT CTTCC GATCT YRYRG TGCCA GCMGC CGCGG TAA-3′ and PE16S-V4-E786-R, 5′- CGGCA TTCCT GCTGA ACCGC TCTTC CGATC TGGAC TACHV GGGTW TCTAA T-3′). Cycle conditions for the first step were: initial denaturation at 98 °C for 30s, denaturation at 98 °C for 30s, annealing at, 52 °C for 30s, and extension at 72 °C for 30 seconds, 19-22 cycles, depending on concentration of 16S rRNA gene copies in the template. 16S rRNA gene copy number was determined by quantitative PCR using the same cycling conditions as above plus the addition of SYBR Green for 40 cycles on a CFX96 Real-time thermocycler (BioRad). The second step PCR reaction adds sample-specific barcodes (Table S3) and Illumina Miseq sequencing adaptors onto the amplicons using the primers (PE-III-PCR-F, 5′-AATGA TACGG CGACC ACCGA GATCT ACACT CTTTC CCTAC ACGAC GCTCT TCCGA TCT-3′; PEIII- PCR-001-096, 5′-CAAGC AGAAG ACGGC ATACG AGATN NNNNN NNNCG GTCTC GGCAT TCCTG CTGAA CCGCT CTTCC GATCT-3′), following the same temperature cycles described above for 9 cycles. The PCR products after both amplification steps were purified using AMPure XP magnetic beads (Beckman Coulter) at a 0.85:1 bead mix to sample ratio. Libraries were randomly assigned to one of two sequencing batches, pooled, and sequenced (150 bp paired end) on an Illumina Miseq sequencer at Johns Hopkins University Genetic Research Core Facility.

For the 2022 samples, 16S rRNA gene library preparation was performed by University of Maryland Institute for Genome Sciences (Baltimore, MD). The V4 region of the 16S rRNA gene was amplified using the facility’s standard universal V4 primer set (515F, 5’-GTGYC AGCMG CCGCG GTAA-3’ and 806R, 5’-GGACT ACNVG GGTWT CTAAT-3’). Samples were sequences on a NextSeq1000 P1 flow cell, 300 bp reads.

### 4.4 16S rRNA gene amplicon sequence data processing

The amplicon reads were processed with Qiime2 2018.8 [49], following a standard analysis pipeline [42]. Briefly, DADA2 [50] was used to quality trim amplicon reads and call amplicon sequence variants. MAFFT [51] and FastTree [52] were used to perform multiple sequence alignment of ASV sequences and generate a rooted phylogenetic tree. Taxonomic assignments were made using Naïve Bayes classifier trained on the 16S rRNA gene V4 region of Greengenes 13_8 99% OTUs [53]. In Qiime2, all samples were rarefied to 94,840 reads and Shannon’s diversity index [54] and weighted UniFrac distance [55] were calculated. Prediction of function were made using the FAPROTAX2 [56, 57]. As run, FAPROTAX2 requires SILVA taxonomy, so ASV taxonomy was re-assigned using Naïve Bayes classifier trained on the V4 region of SILVA138.2_SSURef_NR99_uniform_classifier_V4-515f-806r [58] for use as an input to FAPROTAX2.

### 4.5 Shotgun metagenome sequencing and analysis

Shotgun metagenomics sequencing was performed on two samples collected from surface waters at RC1 and RC2 on 7/23/2019, during the peak of the bloom. DNA was extracted from filtered water samples as described above. Library preparation was performed using Illumina Nextera XT DNA Library Preparation Kit and sequenced on an Illumina MiSeq system by the Johns Hopkins Genetic Resources Core Facility. Metagenomic data was processed using a standard pipeline [42]. Briefly, reads were trimmed and quality control checked using Trimmomatic [59] and FastQC [60]. Samples with fewer than 100,000 reads were removed from the analysis. Contigs were assembled using SPAdes [61], reads were mapped to contigs using BowTie2 [26], genes calls were made using [62].

### 4.6 Data analysis in R

R version 4.5.1 was used for data analysis and figure generation. Seawater densities were calculated using the UNESCO method in the “oce” package [63]. The “Vegan” package to calculate microbial community diversity metrics based on 16S rRNA gene sequence data [64]. ASV abundance modules were generated using the Weighted Gene Correlation Network Analysis (“WGCNA”) package was used to cluster ASVs into modules of correlated abundance [40]. Though WGCNA was developed to identify gene expression patterns between treatments but has been applied to 16S rRNA gene abundance [65] and environmental time series analysis [66]. WGCNA ASV modules were classified into super-modules using a feature-based rule system. Briefly, for each module the time period of its maximum value was identified as well as any broader temporal trends: increasing, decreasing, “U” shaped trend, or inverted “U” shaped. Based on the shape of the trend, the time period of peak value, and the prominence of the peak, the module was assigned to one of 5 super-modules: “pre-disturbance”, “off”, “altered”, “post-disturbance”, “recovery”, and “no-trend”.

### 4.7 Quality control

To determine the contamination from reagents and other materials, blank samples, created by flushing the tubing with sterile deionized water at the beginning of each sampling trip, were processed with the other samples. Quantitative PCR was performed of samples and blanks using the same reagents, primers, and protocols as were used for step one on the 16S rRNA gene library preparation. Quantification cycle values (Cq value) were significantly lower in samples (18.0 ± 0.21 95%CI) as compared with blanks (24.7 ± 1.21 95%CI), indicating minimal contamination in samples (Table S4).

A total of 144 samples were sequenced which included 115 field samples used in primary analysis as well as, positive controls, seven negative controls and a subsample from randomly selected duplicates (Table S3, Fig. S7). All environmental samples were randomly distributed between two sequencing runs to minimize batch effects. To evaluate reproducibility between the two sequencing runs, five environmental samples and the positive control were sequenced on both runs. Replicate samples between the two sequencing runs had a substantially smaller Weighted UniFrac dissimilarity than samples collected at different stations, times, or depths (Fig. S7), indicating high reproducibility between runs. Negative controls either had a low number of reads (3466 ± 1357) or clustered distantly from environmental samples (Fig. S7), indicating the minimal effect of contamination on data. We compared the microbial composition of the positive control between the theoretical values provided by the manufacturer (Zymo Microbial Community Standard, Zymo Research), the values generated by blasting the ASVs against known 16S rRNA gene sequences of the species included in the positive control, and the values generated by classifying using Greengenes database (the same pipeline used as environmental samples). The results show that the microbial composition generated by Greengenes classification was similar to the one generated by BLAST, except that the genera under the family *Enterobacteriaceae* (*Salmonella* and *Escherichia*) could not be classified to the genus level (Fig. S7). This suggests that the classification step should have captured most of the taxonomic information in the samples, although its resolution for some ASVs might be poor below the family level. Overall, the observed abundances closely matched theoretical abundances.

## Source code and data availability

The raw sequence was deposited at NCBI GenBank sequence read archive BioProject number PRJNA869137. Scripts used to perform trimming, quality control, assembly, alignment, and analysis are stored in the GitHub repository: https://github.com/aliceturnham/RockCreek-Shotgun-Metagenomics-Analysis-2/tree/AT_Edits and https://github.com/GaryZhangYue/RockCreek_16S_2022.

## Acknowledgements

This work was supported by NSF award number 2037775, the JHU Catalyst award given to S.P.P, and NSF CBET-1706416 (L.H., A.H., J.T., L.L.L.). This is UMCES Contribution number XXXX and CBL Ref. No. XXXX-XXX.

